# Prediction of Transcription Factor DNA Binding Affinity with High-Throughput *K*_d_ Measurements and Deep Learning

**DOI:** 10.64898/2026.05.18.725930

**Authors:** Zhi Wang, Di Wang, Ke Shen, Junchen Luo, Xinyao Wang, Nan Wu, Yunzhi Lang, Xiangyu Wang, Jun Ren, Wenyang Dong, Lu Pan, Yitong Lyu, Gang Li, Dubai Li, Chen Xie, Zhen Zhang, Shijun Yu, Liuying Shan, Nannan Zhang, Jian Yan, Mingchen Chen, Xiaoliang Sunney Xie

## Abstract

Transcription factors (TFs) regulate gene expression through specific interactions with genomic DNA. While TF binding motifs from public databases describe sequence preferences, quantifying genome-wide affinity (*K*_d_) is highly desirable for a more accurate thermodynamic description. Here, we report *ivt*FOODIE (*in vitro* FOOtprinting with DeamInasE), an assay that leverages deaminase-mediated cytosine-to-uracil conversion to measure *K*_d_ values for a given TF across accessible genomic regions from human cells. By pre-training on TF binding sites from JASPAR and fine-tuning with our *ivt*FOODIE data from 46 TFs representing 13 different DNA-binding domains (DBDs), we developed Seq2K_d_, a deep learning model capable of predicting a TF’s absolute binding affinity on DNA sequences. Seq2K_d_ enables *de novo* motif discovery of ∼500 previously uncharacterized human TFs and reveals the effects of genetic variation both in TF-coding regions and DNA-binding sites on gene expression and disease susceptibility. By correlating predicted affinity changes with the sign and magnitude of expression quantitative trait locus (eQTL) effects, we stratified TFs into activator-like and repressor-like groups. Compared to clinically benign variants, pathogenic single-nucleotide variants (SNVs) within regulatory and protein-coding regions show significantly larger predicted shifts in *K*_d_. We provide an interactive web portal, the ENcyclopedia of Transcription-factor Interactions with Regulatory Elements (ENTIRE), which integrates the Seq2K_d_ model with the *ivt*FOODIE dataset. This resource offers thermodynamic prediction for TF–DNA interactions for functional genomics and human disease.

## Main Text

Transcription factors (TFs) orchestrate gene expression through sequence-specific interactions with genomic DNA (*1*). A central challenge in functional genomics is to quantitatively map the regulatory logic of TF–DNA interactions. This logic is dictated by both sequence specificity, which defines binding locations, and absolute binding affinity, which determines the thermodynamic stability of TF–DNA complexes.

TF–DNA binding specificity has been described primarily through the concept of a binding motif, which is derived from a collection of short homologous nucleotide sequences. These motifs are classically represented by position weight matrices (PWMs), which can be visualized as sequence logos (*2*). Typically, a binding motif can be determined experimentally from observed bound sequences using methods such as Chromatin ImmunoPrecipitation (ChIP-chip (*3*) and ChIP-seq (*4*)), Protein-Binding Microarrays (PBMs) (*5*) and High-Throughput Systematic Evolution of Ligands by EXponential enrichment (HT-SELEX) (*6*), leading to the establishment of large open-access databases such as JASPAR (*7*) and HOCOMOCO (*8*).

For a particular TF on a DNA sequence, the PWM-based motif provides a statistical proxy for sequence preference, whereas the TF’s *K*_d_ provides a direct physical measurement that spans several orders of magnitude. Classic biochemical methods for measuring TF–DNA *K*_d_, such as electrophoretic mobility shift assays (EMSAs) (*9*) and surface plasmon resonance (SPR) (*10*), are inherently low-throughput. Although scalable strategies, including Mechanically Induced Trapping Of Molecular Interactions (MITOMI) (*11*) and High-Throughput Sequencing-Fluorescent Ligand Interaction Profiling (HiTS-FLIP) (*12*), have generated extensive binding data using synthetic DNA libraries, they rely on specialized instrumentation and complex experimental workflows. Thus, direct, simple, and reliable quantification of *K*_d_ within native sequence contexts remains challenging.

Computationally, pioneering deep-learning models (e.g., DeepSEA (*13*), DeepBind (*14*)) excel at predicting categorical binding sites but lack thermodynamic quantification. While ProBound (*15*) predicts relative binding strengths, it cannot generalize to uncharacterized TFs. Notably, AlphaFold3 predicts static structures for given TF–DNA complexes but does not inherently provide thermodynamic affinity data (*16*).

To overcome the above limitations, we report *ivt*FOODIE (*in vitro* FOOtprinting with DeamInasE), a high-throughput sequencing-based *in vitro* approach for genome-wide TF *K*_d_ measurements using double-stranded DNA (dsDNA) cytosine deaminases (*17*). We used high-quality *ivt*FOODIE *K*_d_ data from 46 TFs to develop Seq2K_d_, a deep learning framework that leverages Evolutionary Scale Modeling 2 (ESM2) (*18*), a protein language model, to predict *K*_d_ values for TF–DNA sequence pairs.

### *ivt*FOODIE measures TF binding affinity across accessible human genomic regions

To determine the genome-wide binding affinities (*K*_d_) of TFs on human cis-regulatory elements (CREs), we established the *ivt*FOODIE workflow (Fig. 1A). Briefly, accessible chromatin regions enriched by Assay for Transposase-Accessible Chromatin sequencing (ATAC-seq) (*19*) were incubated with a twofold serial dilution series of the purified target TF spanning the micromolar-to-nanomolar range. Upon reaching binding equilibrium, the high-efficiency dsDNA cytosine deaminase MGYPDa829 (*20*) was introduced for a limited time to catalyze cytosine-to-uracil conversions at unbound sites (Table S1 and Methods). Deep sequencing and conversion-ratio analyses of these loci enabled the *de novo* identification of TF footprints at near-single-base resolution and their fractional occupancy across varying TF concentrations (Fig. S4, B to D and Methods). Site-specific *K*_d_ values were subsequently derived by fitting the Hill equation to these occupancy data (Fig. S1, C to F and Methods). The ATAC-seq-enriched open chromatin from GM12878 cells used as input for *ivt*FOODIE libraries faithfully recapitulated open chromatin profiles (Fig. S1, A and B). *ivt*FOODIE facilitates high-resolution footprinting and quantitative affinity measurements, capturing thermodynamic variations across individual TF footprints with high throughput (Fig. 1B and Fig. S4E).

**Fig. 1.**
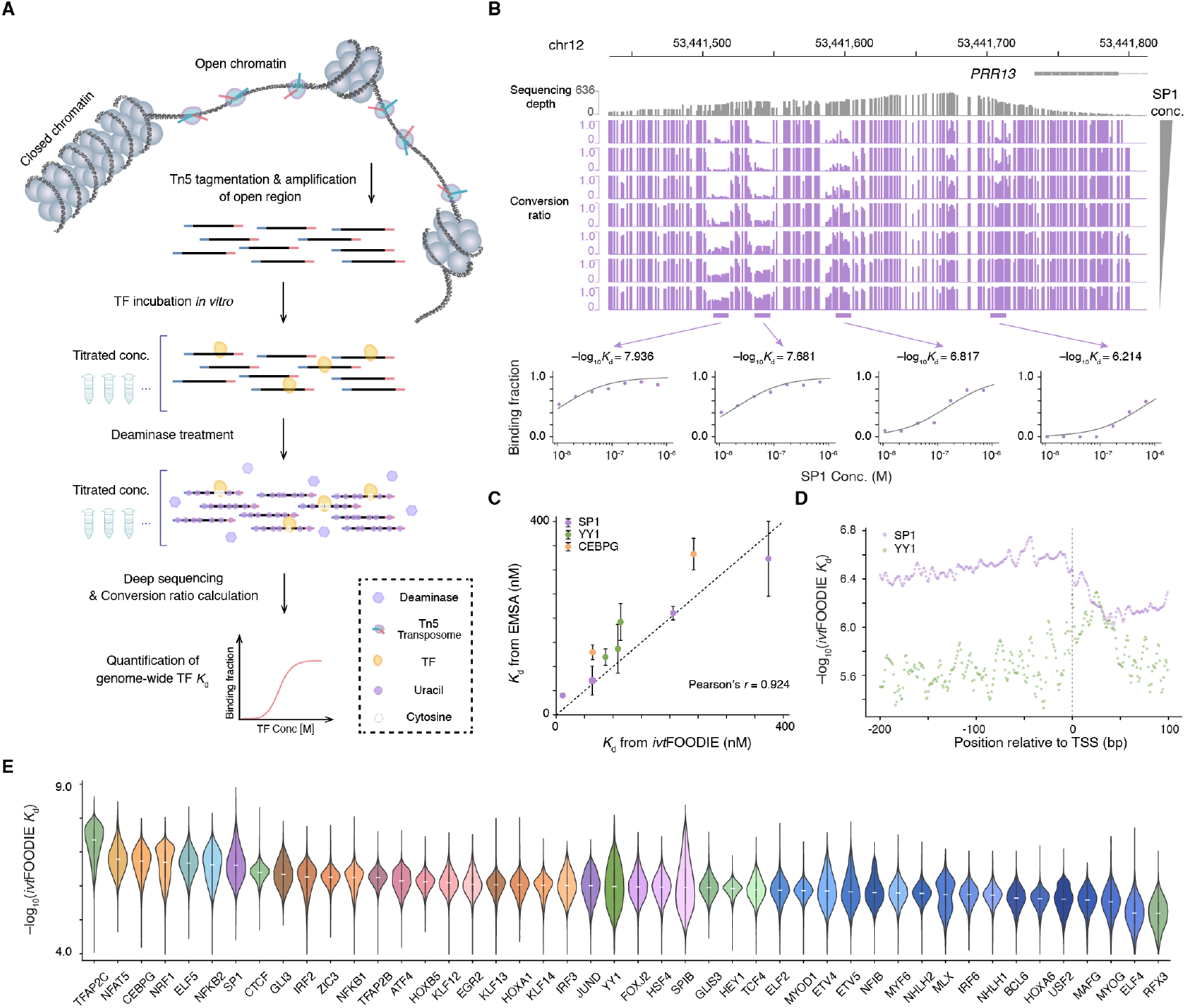
*ivt*FOODIE enables high-throughput detection of binding affinities across accessible human genomic regions. **(A)** Schematic of *ivt*FOODIE. Open chromatin DNA from GM12878 cells is prepared through Tn5 tagmentation, PCR amplification, and size selection. A TF of interest is incubated with the DNA at varying concentrations until thermodynamic equilibrium is reached. Binding fractions across the genome are quantified based on C-to-U conversion rates after deaminase treatment, followed by deep sequencing. Site-specific binding affinities are derived by fitting the binding fractions observed at different TF concentrations. **(B)** SP1 binding at the *PRR13* promoter. Top: sequencing depth (gray) and C-to-U conversion signals (purple) across an SP1 concentration gradient (right wedge). Four discrete SP1 binding sites (purple bars) show decreased conversion (increased protection) at higher SP1 concentrations. Bottom: fitted binding curves (solid lines) and observed fractions (circles) used to calculate *K*_d_. **(C)** Comparison of *K*_d_ measured by EMSA and *ivt*FOODIE. Each dot represents one sequence bound by a given TF, and error bars indicate the standard deviation from three independent measurements. **(D)** Differential TF binding affinity preferences around TSSs. Mean –log_10_*K*_d_ is plotted as a function of genomic distance from the TSS (position 0). Data were smoothed using a 3-bp Gaussian kernel. **(E)** Violin plots of – log_10_*K*_d_ for 46 TFs measured in GM12878 cells, ordered by median affinity. White horizontal bars denote medians, and vertical black lines represent the full range of the data (filtered for *–*log_10_*K*_d_ > 4.0).

Orthogonal validation experiments using EMSA for SP1, YY1, and CEBPG demonstrated strong quantitative concordance with *ivt*FOODIE-derived *K*_d_ measurements across *de novo* footprints (Fig. 1C, Fig. S2, A to I and Table S2). Benchmarking *ivt*FOODIE-derived *K*_d_ measurements against enrichment scores from published HT-SELEX datasets also demonstrated high concordance (Fig. S3A).

Global *K*_d_ profiles revealed distinct spatial preferences. YY1 exhibited enhanced affinity downstream of transcription start sites (TSSs), while SP1 displayed a marked preference for upstream regions (Fig. 1D and Fig. S3B). These *in vitro* binding landscapes are highly concordant with *in vivo* ChIP-seq results (Fig. S3C) and align with prior motif-based observations (*21*).

Notably, TFs exhibit distinct thermodynamic behaviors across different genomic contexts. For instance, while certain factors such as SP1 and various KLF members (e.g., KLF12, KLF13, KLF14) maintain comparable binding affinities between promoters and enhancers, others show a marked preference for specific elements. Specifically, consistent with their known binding preferences and DNA sequence distribution, *ivt*FOODIE data show that NRF1 and YY1 bind to promoter regions with significantly higher affinity than to enhancers (Fig. S4, F and G). These results suggest that TFs fine-tune their regulatory functions by using differential *K*_d_ across CREs.

Expanding this approach, we generated genome-wide *K*_d_ landscapes for 46 human TFs (Table S1). The profiles revealed that affinity distributions varied in both magnitude and breadth across different TF families (Fig. 1E). At one extreme, TFAP2C from the AP-2 TF family exhibited a broad range of high affinities (median –log_10_*K*_d_ > 8.0), reflecting stringent base readout and rigid protein–DNA interfaces. Conversely, the architectural protein CTCF exhibited a narrow affinity distribution range, indicating uniform binding strength across the genome. Unlike typical TFs that require variable affinities to fine-tune gene expression, CTCF relies on its multiple zinc fingers to strictly recognize target sequences. This uniform and stable binding is consistent with CTCF’s architectural role in chromatin organization.

### Seq2K_d_ predicts TF affinity on DNA sequences

Although *ivt*FOODIE provides high-fidelity measurements, experimentally profiling all ∼1,600 human TFs remains impractical due to protein purification constraints. Given that the vast majority of TFs cluster into a limited number of structural DNA-binding domain (DBD) families (∼30), we reasoned that a deep learning framework could learn a generalizable biophysical grammar from a sparse experimental matrix. To this end, we utilized JASPAR data for pre-training (Fig. S5, A to D) and *ivt*FOODIE data for fine-tuning.

We present Seq2K_d_, a cross-modal transformer that predicts whether a TF binds to a given DNA sequence (binding classification), the quantitative *K*_d_ value for bound pairs, and the corresponding prediction error (pError). Protein sequences are encoded by ESM2 and DNA by a dedicated encoder, and the two are fused through cross-modal attention for bidirectional information exchange. A shared multi-task head simultaneously optimizes binding classification and *K*_d_ regression via a composite loss. Training involves three stages: (i) masked nucleotide recovery pre-training to learn TF–DNA grammar from JASPAR binding sequences; (ii) joint fine-tuning for binding classification, *K*_d_ prediction, and error estimation using *ivt*FOODIE *K*_d_ data; (iii) self-distillation using augmented TF–DNA pairs to improve calibration of predicted affinities and generalization of binding predictions (Fig. 2A). This final self-distillation step acts as a data augmentation strategy that uses virtual data to expand the learning space and enhance predictive accuracy. This unified architecture and training pipeline integrate heterogeneous data for accurate absolute *K*_d_ prediction for TF–DNA pairs.

**Fig. 2.**
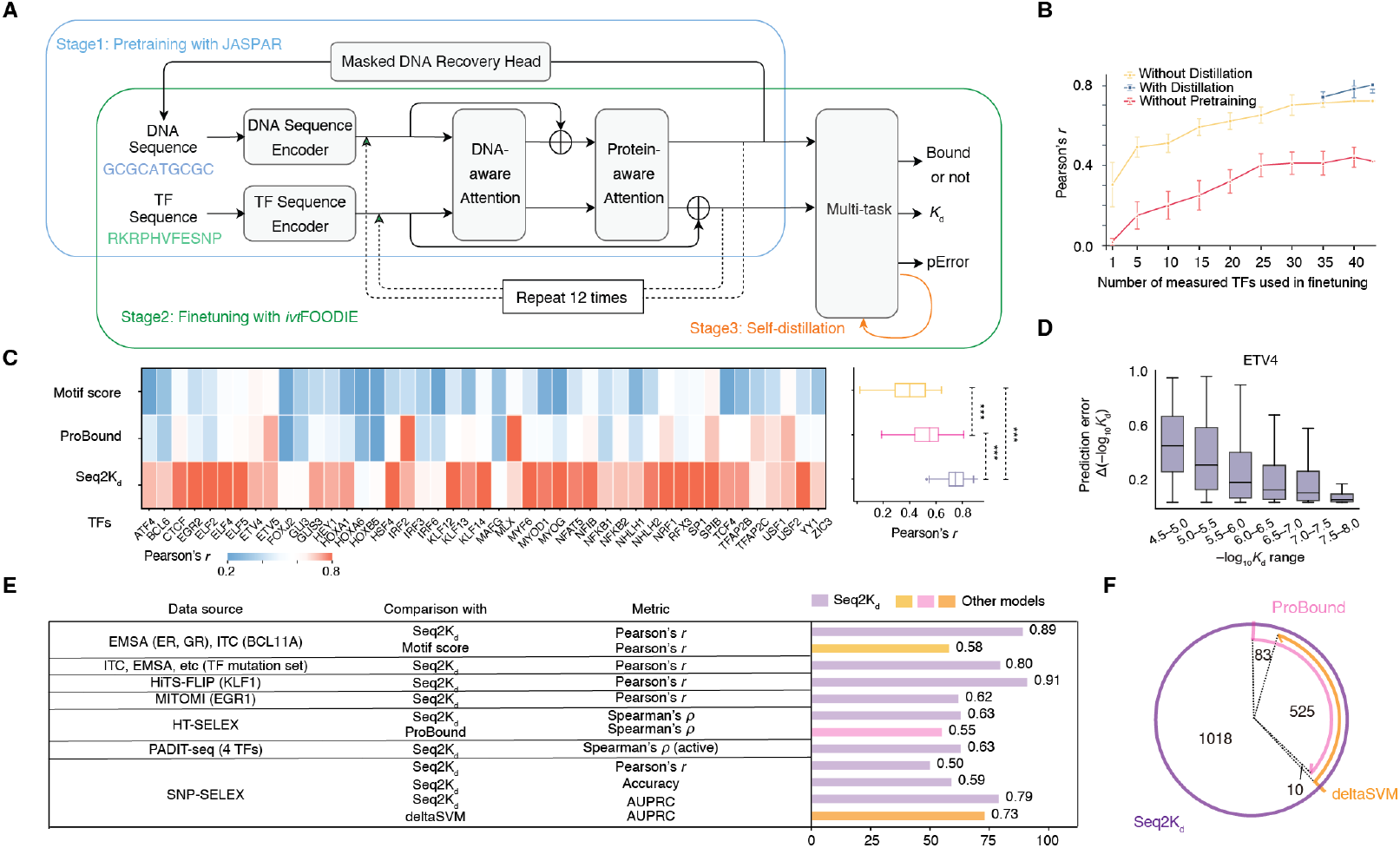
Architecture, training, and performance evaluation of the Seq2K_d_ model. **(A)** Seq2K_d_ architecture and training strategy. The model is trained via a three-stage pipeline: pre-training on JASPAR TF-binding data, fine-tuning with *ivt*FOODIE *K*_d_ values, and self-distillation. The core architecture features a 12-layer DNA–protein attention module, culminating in a multi-task head that simultaneously predicts binary binding probability, absolute affinity (*K*_d_), and the *K*_d_ prediction error (pError). **(B)** Scaling laws for TF generalization. Pearson’s *r* between *ivt*FOODIE-measured *K*_d_ values and predictions from Seq2K_d_, Seq2K_d_ without pre-training, and Seq2K_d_ without distillation for three held-out TFs evaluated across increasing training set sizes. The self-distilled Seq2K_d_ model consistently outperforms the non-distilled baseline. **(C)** Leave-one-TF-out cross-validation. Pearson’s *r* across individual held-out TFs (heatmap, left). Boxplots summarize the results (right), demonstrating that Seq2K_d_ achieves superior predictive correlation. Center line, median; box limits, upper and lower quartiles; whiskers, 1.5× interquartile range. Asterisks denote statistical significance (*P* < 0.001, two-sided Wilcoxon signed-rank test). **(D)** Relationship between prediction error and affinity. For the representative TF ETV4, high-affinity interactions (larger –log_10_*K*_d_) consistently exhibit lower pError, indicating higher confidence in strong binding predictions. **(E)** Benchmarking of Seq2K_d_ against established computational methods using various independent *in vitro* datasets. The table outlines the data sources (including EMSA (*22-26*), ITC (*27-30*), Fluorescence Anisotropy (FA) (*41-43*), Fluorescence Polarization (FP) (*44*), MicroScale Thermophoresis (MST) (*45*), BioLayer Interferometry (BLI) (*46, 47*), HiTS-FLIP (*31*), MITOMI (*32*), HT-SELEX (*6*), PADIT-seq (*33*), and SNP-SELEX (*34*)) and the specific statistical metrics used for evaluation (Pearson’s *r*, Spearman’s *ρ*, accuracy, and AUPRC (Area Under the Precision–Recall Curve)). In the adjacent bar plot, purple bars indicate the performance of Seq2K_d_, while the yellow, pink, and orange bars denote the performance of the respective comparison models (motif score, ProBound, and deltaSVM). Exact performance scores are annotated at the end of each bar. **(F)** Expanded prediction coverage across the human TF repertoire. Proportional Euler diagram illustrating the number of human TFs whose binding affinities can be modeled by Seq2K_d_ (purple), deltaSVM (orange), and ProBound (pink). Seq2K_d_ substantially expands the predictable TF repertoire.

Model performance plateaued upon fine-tuning with *ivt*FOODIE data for ∼40 TFs, achieving a Pearson’s *r* of ∼0.8 (Fig. 2B). Systematic component analysis confirmed the critical roles of our key architectural designs in achieving high accuracy: multi-task learning, cross-modal attention, and self-distillation (Methods and Supplementary Information). Notably, while self-distillation yielded marginal gains, large-scale pre-training proved indispensable, driving a substantial performance improvement (Δ*r* ≈ 0.3). Model interpretability analyses revealed that the learned attention maps correspond closely with experimental protein–DNA contacts (Supplementary Information).

To evaluate unseen TFs at the individual level, we systematically excluded each TF from the *ivt*FOODIE dataset during Seq2K_d_ training. Evaluations of these held-out TFs revealed that Seq2K_d_ consistently outperformed both PWM motif scores and ProBound in correlating with experimentally measured *K*_d_ values (Fig. 2C). We further observed a strong correlation between prediction error and binding affinity. Specifically, across four held-out TFs (CTCF, ELF2, ELF4, and HSF4), high-affinity DNA sequences, which have lower *K*_d_ values, consistently exhibited reduced prediction errors (Fig. 2D and Fig. S5E). To assess generalizability, we evaluated Seq2K_d_ on held-out TF clusters sharing less than 70% DBD sequence identity with the training set. Across these independent test cohorts, Seq2K_d_ maintained predictive accuracy (average Pearson’s *r* ≈ 0.6) (Supplementary Information).

Moreover, Seq2K_d_ consistently performed better than existing models across a wide range of independent datasets (Fig. 2E, Fig. S6 and Fig. S7). Across traditional biophysical assays (EMSA (*22-26*), Isothermal Titration Calorimetry (ITC) (*27-30*), etc.), modern high-throughput methods (HiTS-FLIP (*31*), MITOMI (*32*), HT-SELEX (*6*), PADIT-seq (*33*), SNP-SELEX (*34*)), and comparisons with existing predictive models (ProBound (*15*) and deltaSVM), Seq2K_d_ provided highly accurate *K*_d_ predictions for key TFs, demonstrating high reliability. Furthermore, Seq2K_d_ is not restricted to a limited subset of TFs and generalizes predictions across the entire human TF repertoire (Fig. 2F).

### *De novo* motif discovery for uncharacterized TFs

While valuable resources such as JASPAR and HOCOMOCO provide motifs for ∼1,100 human TFs, the binding specificities of ∼500 (out of ∼1,600) remain experimentally uncharacterized (Fig. 3A). Furthermore, because individual members of the same TF family can possess highly distinct core motifs, we harnessed the generalizability of Seq2K_d_ to capture this diversity. We leveraged Seq2K_d_ to compute high-affinity genomic sites and derive *de novo* motifs for these uncharacterized TFs, for which experimental characterization has been challenging (Fig. S8, A to C and Methods).

**Fig. 3.**
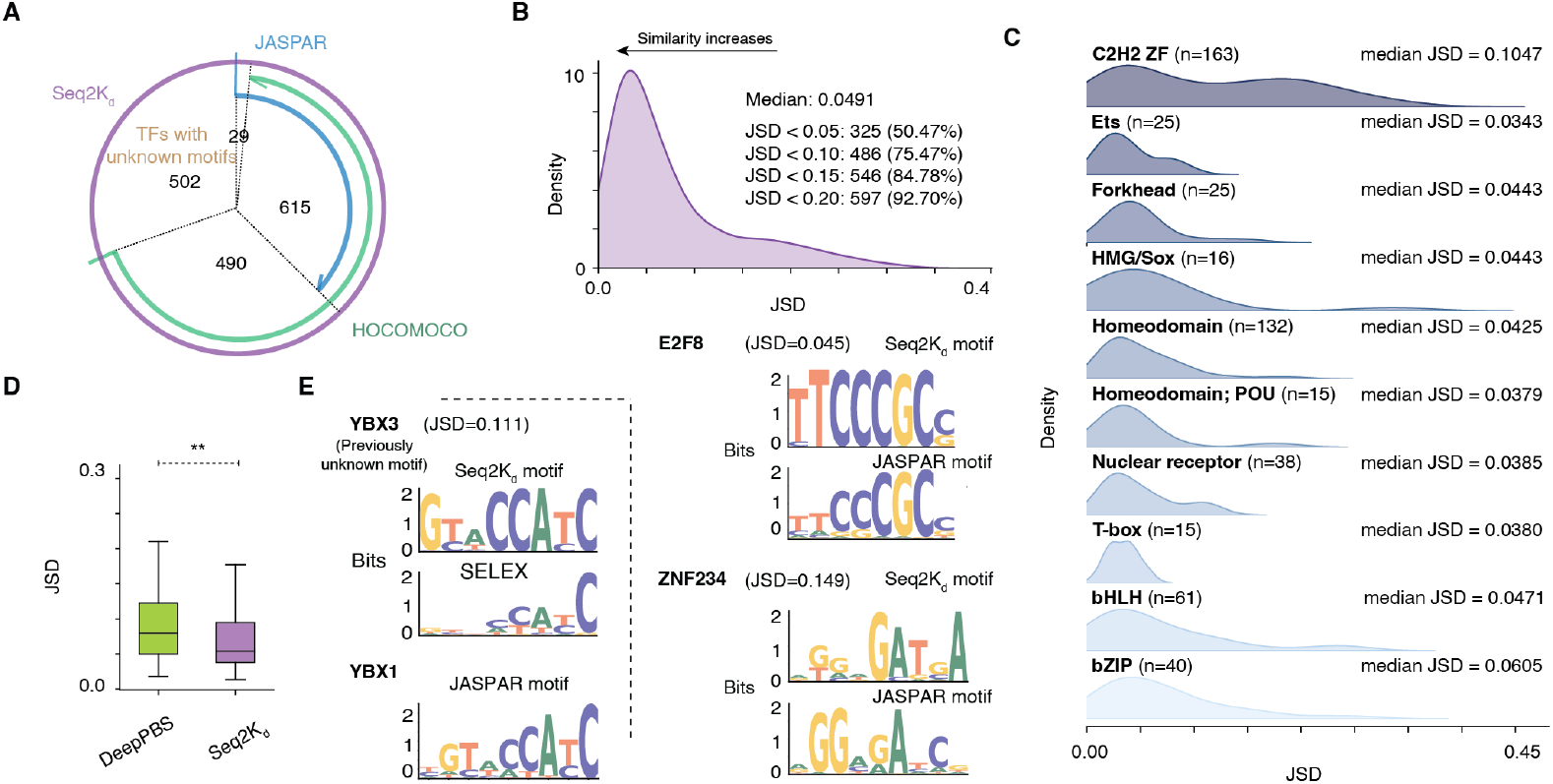
Seq2K_d_ accurately derives TF binding motifs *de novo*. (**A)** Expanded motif coverage across the human TF repertoire. Proportional Euler diagram comparing the number of human TFs with binding motifs derivable by Seq2K_d_ (purple) versus those with reference motifs in the JASPAR (blue) and HOCOMOCO (green) databases. Seq2K_d_ derives motifs for 502 TFs identified by Lambert et al. (*1*) that previously lacked binding specificity. **(B)** Global accuracy of predicted motifs. Top: Density plot of the JSD between Seq2K_d_-predicted motifs and JASPAR references. Inset text denotes the median JSD and the cumulative percentage of TFs below specific JSD thresholds. Bottom: Comparisons of Seq2K_d_-predicted motifs against known references. E2F8 and ZNF234 illustrate cases of small and large JSD, respectively. **(C)** Seq2K_d_–JASPAR motif concordance across TF families. Density distributions of JSD scores stratified by major TF structural families and visualized using a gradient of blue hues. *n* represents the number of TFs per family; family-specific median JSD scores are annotated. **(D)** Benchmark of motif prediction. Box plots comparing JSD scores of JASPAR motifs to motifs derived by DeepPBS and Seq2K_d_. Asterisks denote statistical significance (*P* < 0.01, two-sided Wilcoxon rank-sum test). **(E)** Validation of *de novo* motif prediction. The Seq2K_d_-derived motif for YBX3 (which lacks a reference) matches its motif determined by SELEX. The JASPAR motif for the homolog YBX1 is shown for comparison. Y-axis: information content (bits).

We used Jensen-Shannon divergence (JSD) to evaluate Seq2K_d_’s accuracy against curated databases. Unlike standard distance metrics, JSD quantifies the informational overlap between probability distributions, thereby serving as a proxy for capturing differences in thermodynamic binding affinities. JSD scores were highly concordant with results from Tomtom, which is widely used to compare a query motif against a database of known motifs (Fig. S8, D and E and Methods). Our predictions exhibited high global concordance with JASPAR TF motifs (median JSD = 0.049), with 75.5% of TFs showing highly similar profiles (JSD < 0.1) (Fig. 3B and Fig. S8G). Accuracy varied by TF structural family (Fig. 3C and Fig. S8H). Families with rigid specificities like ETS (*n* = 25) exhibited minimal divergence (median JSD = 0.034), whereas C2H2 zinc-finger (C2H2-ZF) TFs (*n* = 163) showed higher deviation (median JSD = 0.105) that correlated positively with the number of zinc-finger domains (Pearson’s *r* = 0.5; Fig. S8J). To assess generalizability, we evaluated Seq2K_d_ on a held-out set of 200 human TFs after excluding proteins with >70% DBD sequence identity to any protein in the training data. This set yielded a lower median JSD than the training set (Fig. S8, J to L), likely due to a lower proportion of challenging C2H2-ZFs (Fig. S8I and Methods).

When compared with JASPAR motifs as the reference, motifs derived by Seq2Kd exhibited a significantly lower median JSD than those derived by the structure-based prediction algorithm DeepPBS (*35*) (Fig. 3D and Fig. S8F). These results indicate that Seq2K_d_ outperforms the DeepPBS framework in *de novo* motif prediction.

For orthogonal experimental validation, we performed a SELEX experiment on YBX3, a TF without any previously characterized motif. Although the entire YBX family was strictly excluded from both pre-training and fine-tuning datasets, the experimentally derived *in vitro* motif strongly matched the Seq2K_d_ *de novo* prediction (Fig. 3E). We used Seq2K_d_ to generate motifs *de novo* for all previously uncharacterized human TFs and compiled these predictions into a new open-access platform, ENTIRE (ENcyclopedia of Transcription-factor Interactions with Regulatory Elements; https://www.entire.ac.cn/), providing a resource for exploring regulatory mechanisms of poorly studied TFs (Fig. S10, A to E).

### Linking mutation-induced Δ*K*_d_ to gene expression

We next examined the *in vivo* relevance of the predicted *K*_d_ values to gene expression. Directly predicting in vivo TF occupancy from *K*_d_ is challenging because local TF concentrations are generally unknown, and multiple TFs may compete for the same or overlapping binding sites. We therefore turned to expression quantitative trait loci (eQTLs), which bypass the need to explicitly model local TF concentration and chromatin competition while providing a direct functional readout of regulatory variant effects.

We then evaluated whether predicted affinity changes (Δ(*–*log_10_*K*_d_), defined as *–*log_10_*K*_d_ of the mutant sequence minus *–*log_10_*K*_d_ of the reference sequence, hereafter Δ*K*_d_) across regulatory single-nucleotide variants (SNVs) could mechanistically explain eQTL effect sizes provided by GTEx (*36*). By correlating Δ*K*_d_ with eQTL effect sizes, we classified human TFs by inferred regulatory directionality, with positive Δ*K*_d_–eQTL correlations indicating activator-like behavior and negative correlations indicating repressor-like behavior (Fig. 4A, Fig. S10, A and B). This stratification was supported by Gene Ontology (GO) enrichment (Fig. 4B). Furthermore, this Δ*K*_d_-based framework outperformed standard ΔPWM scoring in recovering TFs annotated as transcriptional activators, using GO annotations as a benchmark.

**Fig. 4.**
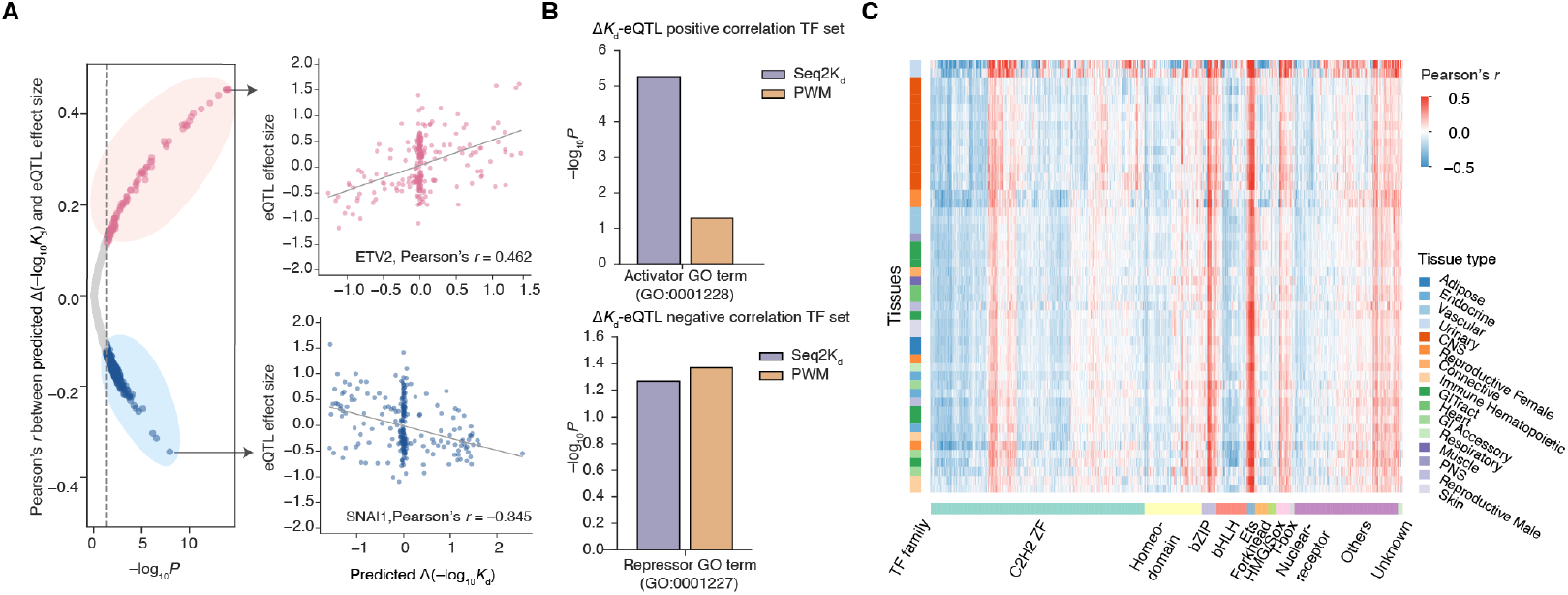
Predicted TF binding affinity changes at promoter TF binding site mutations correlate with gene expression. **(A)** Association between predicted affinity changes (Δ(*–*log_10_*K*_d_), hereafter Δ*K*_d_) and eQTL effect size across the genome, where each dot represents a single human TF. Pink and blue shaded regions highlight TFs with significant positive or negative associations, respectively. Scatter plots to the right show predicted Δ*K*_d_ (x-axis) versus eQTL effect size (y-axis) for representative TFs showing positive (ETV2, top) and negative (SNAI1, bottom) correlations. **(B)** Gene Ontology (GO) functional enrichment of eQTL-associated TFs. TFs exhibiting significant positive (top) or negative (bottom) correlations between Δ*K*_d_ and eQTL effect size are enriched for activator (GO:0001228) and repressor (GO:0001227) functions, respectively. Bar heights indicate *–*log_10_*P* values for enrichments using Seq2K_d_ (purple) or PWM (orange) models. **(C)** Δ*K*_d_–eQTL correlations across tissues. Heatmap displaying Pearson’s *r* between predicted Δ*K*_d_ and eQTL effect sizes across different tissue types (rows) for various TFs (columns, clustered by family). Red and blue indicate positive and negative correlations, respectively.

Extending this classification to GTEx tissues revealed that the Δ*K*_d_–eQTL correlations remain broadly consistent across multiple tissues, albeit with subtle tissue-dependent variations (Fig. 4C). For instance, the bZIP and ETS families consistently displayed strong activator-like behavior across various tissues, aligning with their established roles in broad transcriptional activation. Furthermore, TFs classified as activator-like in most tissues showed stronger activator-like signatures in urinary system-associated tissues.

### Linking mutation-induced Δ*K*_d_ to human disease

To investigate the relationship between TF binding perturbations and disease, we evaluated variants from ClinVar (*37*). Across both promoter SNVs and TF-coding mutations, pathogenic variants exhibited significantly greater *K*_d_ perturbations than clinically benign variants (Fig. 5A and Fig. S10, C and D), supporting an association between larger predicted affinity perturbations and pathogenic classification.

**Fig. 5.**
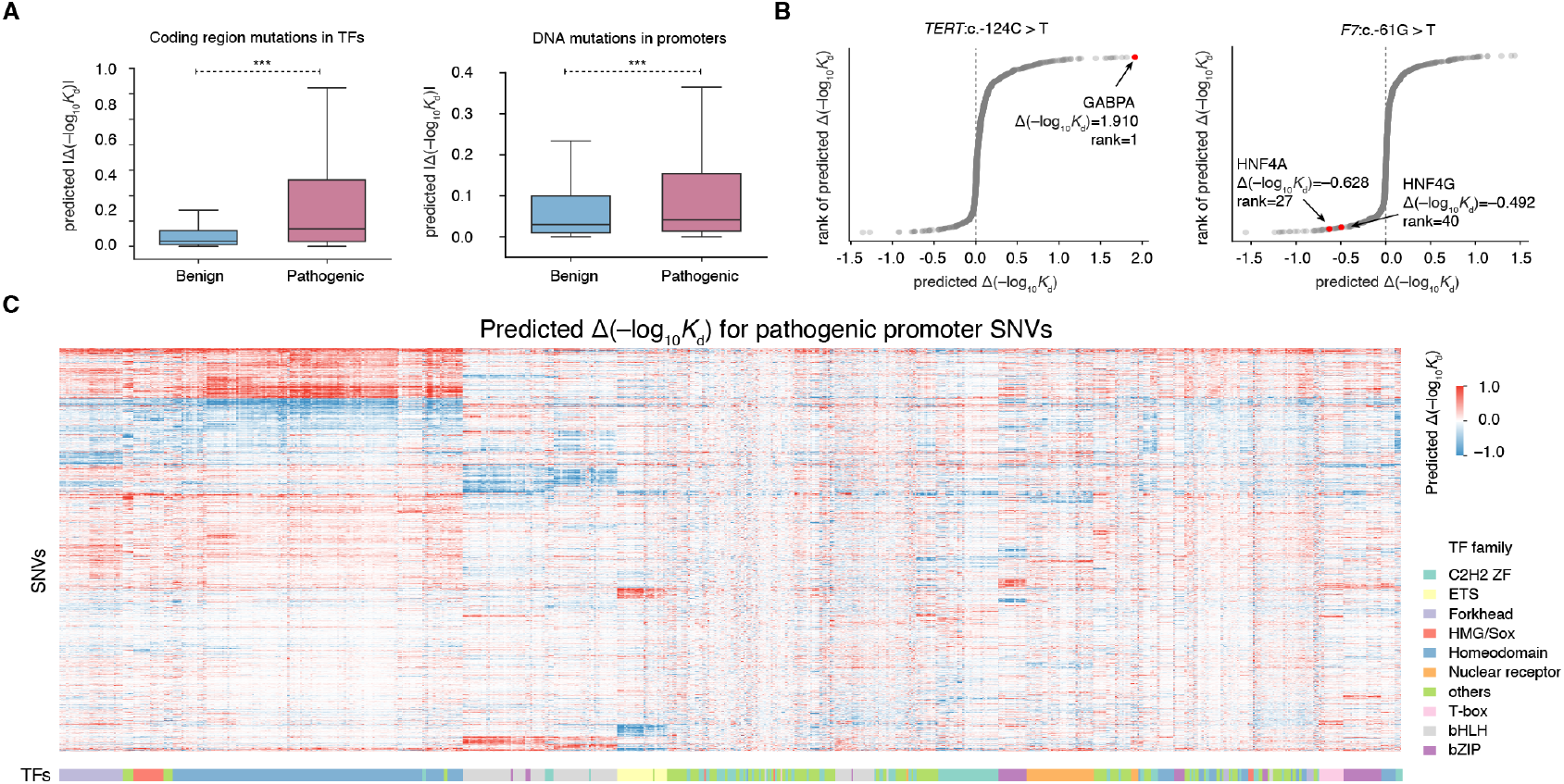
Predicted TF binding affinity changes associated with human disease variants. **(A)** Impact of variants on affinity by pathogenicity. Boxplots comparing the absolute magnitude of predicted Δ*K*_d_ induced by benign (blue) versus pathogenic (pink) variants curated in ClinVar. Variants are categorized as coding region mutations in TFs (left) or DNA mutations in promoters (right). Center lines indicate medians, box bounds represent the interquartile range (IQR), and whiskers extend to 1.5 × IQR. \*\*\**P* < 0.001 (two-sided Wilcoxon rank-sum test). **(B)** Identification of specific pathogenic non-coding SNVs. Left, scatter plots showing the rank versus signed value of predicted Δ*K*_d_ across all human TFs for the disease-associated promoter mutations *TERT* c.-124C>T (left, associated with cancer) and *F7* c.-61G>T (right, associated with Factor VII deficiency). Arrows highlight the experimentally-confirmed TFs, GABPA and HNF4A/HNF4G, respectively. **(C)** Predicted Δ*K*_d_ landscape for pathogenic promoter variants in ClinVar. Heatmap visualizing the predicted Δ*K*_d_ induced by curated ClinVar promoter SNVs (rows) across human TFs clustered by family. Red indicates increased binding affinity, and blue indicates decreased affinity.

Beyond these cohort-level statistics, Seq2K_d_ captured significant affinity shifts on well-characterized, disease-causing non-coding mutations. For instance, the model accurately predicted the previously known pathogenic affinity gain for GABPA at the oncogenic *TERT* promoter mutation (c.-124C>T) (*38*), as well as the known severe affinity loss for HNF4 family members, including HNF4A and HNF4G, at the monogenic *F7* promoter variant (c.-61G>T) that causes hereditary Factor VII deficiency (*39*) (Fig. 5B). Additional case studies reliably recovered established TF-variant associations across metabolic and immune disorders (*40*) (Fig. S10E).

Motivated by the model’s accuracy on these established clinical cases, we systematically profiled all SNVs in ClinVar to construct a resource of TF affinity disruptions (Fig. 5C, Fig. S10D). Analysis of this global resource revealed that the pathogenic disruptions disproportionately affected TF families associated with master developmental regulators, such as the homeodomain, SOX, and GATA families, suggesting that changes in their thermodynamic binding characteristics are associated with disease. Collectively, these results establish that integrating thermodynamic affinity modeling with population and clinical genetics provides a unified framework to decode how regulatory variants drive human disease.

## Discussion

PWMs are widely used proxies for TF–DNA binding, but their pairwise additive formulation does not account for dependencies between nucleotide positions. Using the *ivt*FOODIE dataset, we directly quantified this limitation: across diverse TFs, Seq2K_d_ correlated more strongly with experimental measurements than conventional PWM-based scores (Fig. 2C), consistent with recent biophysical studies *(48)*. These results indicate that Seq2K_d_, by capturing sequence dependencies beyond additive motif representations, can improve the description of TF–DNA binding specificity.

Unlike synthetic library assays, *ivt*FOODIE interrogates binding in the context of native genomic DNA. Although its reliance on cytosine deamination theoretically introduces a compositional bias against AT-rich regions, Seq2K_d_ maintained high predictive accuracy for AT-rich homeobox TFs (Fig. 3C). This resilience stems from our cross-modal architecture: pan-species JASPAR pre-training establishes a generalized base-recognition grammar (Supplementary Information), which is subsequently calibrated by high-resolution *ivt*FOODIE regression.

Despite originating from the accessible genome of a single cell line (GM12878), the *ivt*FOODIE dataset encompasses sufficient genomic sequence diversity to train Seq2K_d_ effectively. Training on only 46 TFs across 13 major families achieved strong performance across the broad spectrum of human TFs, with room for improvement among some zinc-finger TFs (Fig. 3C). Extending *ivt*FOODIE measurements to additional TF families will further refine these models.

Although currently fine-tuned on human *ivt*FOODIE profiles, the cross-species pre-training of Seq2K_d_ on the JASPAR dataset—spanning mouse, *Drosophila, C. elegans*, and yeast—positions it to generalize across diverse taxa. Complementing this, the integration of the ESM2 protein language model allows Seq2K_d_ to capture how evolution shapes the underlying biophysics of DNA recognition, empowering the model to assess non-canonical DNA-binding proteins and uncharacterized regulators.

Overall, the *ivt*FOODIE assay establishes a high-throughput, single-base-resolution thermodynamic baseline for the genome-wide landscape of TF–DNA binding affinity. By distilling these extensive measurements through deep learning, Seq2K_d_ predicts absolute binding affinities across the entire human TF repertoire, creating a comprehensive map of regulatory interactions. These predicted affinities provide a mechanistic bridge linking genetic variants to both gene expression modulation and clinical pathogenicity. We provide this unified resource via the publicly available ENTIRE platform, offering a quantitative framework for decoding functional genomics and disease mechanisms.

## Acknowledgments

This project was financially supported by Changping Laboratory and the Ministry of Science and Technology of China. Xinyao W was supported in part by the Peking University Boya Postdoctoral Fellowship. We are grateful to Prof. Jussi Taipale (Karolinska Institute) for providing the cDNA collection of several human TFs.

## Funding

Changping Laboratory (2021C0402)

Shenzhen Medical Research Fund (B2302027 and D2501003)

Research Grants Council of HKSAR (11101022)

National Natural Science Foundation of China (32270634)

## Author contributions

Conceptualization: XSX, MC, ZW

*ivt*FOODIE methodology: ZW, Xinyao W, SY, WD (deaminase purification), XSX

Data collection: ZW, Xinyao W, SY, JR, LS, NZ, NW (TF protein purification), YL (SELEX experiments)

Seq2K_d_ algorithm development: MC, DW, KS, JL

Data analysis: KS, DW, JL, Xiangyu W, YL, ZW, GL, DL, CX, ZZ

Visualization: ZW, KS, DW, JL, Xinyao W, Xiangyu W, YL

Website construction: LP, DW, YL

Funding acquisition: XSX, MC, JY

Supervision: XSX, MC, JY

Writing: XSX, MC, ZW, KS, JL, YL, Xinyao W, Xiangyu W, with input from all other authors.

## Competing interests

XSX, ZW, KS, Xinyao W, and WD are inventors on a patent filed by Changping Laboratory covering the *ivt*FOODIE method described in this work.

