## Supplementary Materials for "Prediction of Transcription Factor DNA Binding Affinity with High-Throughput *K*_d_ Measurements and Deep Learning"

**This Supplementary Materials file includes:**

Materials and Methods
Figs. S1 to S10
tables S1 to S2

### Materials and Methods

#### Purification of TF proteins

The names, full amino acid sequences, and incorporated tags of all TFs used in the *ivt*FOODIE assay are listed in **table S1**. Commercial TFs were obtained from Active Motif. Forty-seven recombinant TFs were expressed using the Gateway system on a modified pETG20A receptor vector, which incorporates an N-terminal thioredoxin-6× His tag and a C-terminal 3× FLAG tag. Three recombinant proteins were expressed using the Gateway system on a modified pET-28a(+) receptor vector, which incorporates an N-terminal thioredoxin-6× His tag and a C-terminal streptavidin-binding peptide (SBP) tag. Expression inserts were derived from the Mammalian Gene Collection (MGC), ORFeome, or via gene synthesis (GenScript).

Recombinant proteins were expressed in *E. coli* and purified following previously described methods *(27)* with minor modifications. Briefly, for ZF proteins, the culture medium was supplemented with 50 µM $ZnSO_4$ to facilitate correct folding. Following cell lysis, the 6× His-tagged proteins were immobilized on Ni-sepharose beads (GE Healthcare, 17-5318-01) in binding buffer (10 mM Tris pH 7.5, 50 mM NaCl, 1 mM $MgCl_2$, 4% glycerol, 0.5 mM EDTA) and eluted after extensive washing. Protein purity was verified by SDS-PAGE and Coomassie brilliant blue staining. Purified proteins were stored at –20°C at a final concentration of 50% (v/v) glycerol.

#### Purification of deaminase MGYPDa829

The genes encoding the deaminase MGYPDa829 (Da829; retaining an N-terminal 6× His tag) and its immunity protein Da829I (S-tag removed) were synthesized and cloned into the MCS-1 and MCS-2 of the pETDuet-1 vector, respectively. Full sequences are provided in **table S1**. Plasmids were maintained in *Escherichia coli* DH5α and transformed into *E. coli* BL21(DE3) for protein expression.

MGYPDa829 was expressed and purified using an on-column denaturation and refolding strategy as previously described *(17)*, with minor modifications. Briefly, following IPTG induction and cell lysis, the 6× His-tagged Da829–Da829I complex was captured using nickel affinity chromatography. The complex was then denatured in 8 M urea to remove Da829I. Bound Da829 was subjected to on-column refolding via a stepwise descending urea gradient and subsequently eluted. The refolded Da829 was further purified by size-exclusion chromatography (Superdex 200, GE Healthcare). Protein purity was assessed by SDS-PAGE.

#### *ivt*FOODIE assay

Open genomic DNA was incubated with TFs in a customized binding buffer at room temperature to facilitate protein–DNA interactions. Following incubation, the complexes were subjected to deamination using the Da829 enzyme. The deamination reaction was quenched via SDS addition and protease digestion, and the converted DNA was column-purified. Finally, sequencing libraries were constructed by amplifying with a uracil-tolerant polymerase (Q5U), and were finalized through bead-based size selection.

#### EMSA

To generate the dsDNA probes for the EMSA, the biotinylated single-stranded oligonucleotides and unlabeled complementary DNA sequences were annealed by heating to 97°C for 5 min, followed by linear cooling to 12°C over 60 min. Single-stranded oligonucleotides were synthesized by Sangon Biotech. The sequences of all biotinylated DNA oligonucleotide templates are detailed in **table S2**. EMSA was performed using the Chemiluminescent EMSA Kit (cat. no. DS009, Beyotime) with minor modifications. For titration experiments, 2 nM biotinylated dsDNA probes were incubated at room temperature for 30 min in a reaction mixture containing commercial or laboratory-purified TFs (twofold serial dilutions ranging from 4 nM to 2 μM), homemade 4× binding buffer (see *ivt*FOODIE Assay above), and 50 ng/μL poly(dI·dC). After reaching thermodynamic equilibrium, the reaction mixtures were separated by 6% native polyacrylamide gel electrophoresis (PAGE) in 0.5× Tris-borate-EDTA (TBE) buffer. Subsequently, samples were transferred to nylon membranes in 0.5× TBE, followed by UV cross-linking and chemiluminescent detection. Signals were captured using the Bio-Rad ChemiDoc Imaging System. Quantitative analysis was performed using ImageJ software (https://imagej.nih.gov/ij). Gel images were first processed by background subtraction, followed by band intensity integration. The fraction of bound DNA (Binding Fraction, BF) was determined from the background-subtracted signal intensities using the following equation:

$$BF = \frac{DNA_{bound}}{DNA_{bound} + DNA_{free}}$$

Binding affinities were calculated by fitting the binding fraction to the sigmoidal equation:

$$BF = \frac{[TF]}{[TF] + K_{\mathrm{d}}}$$

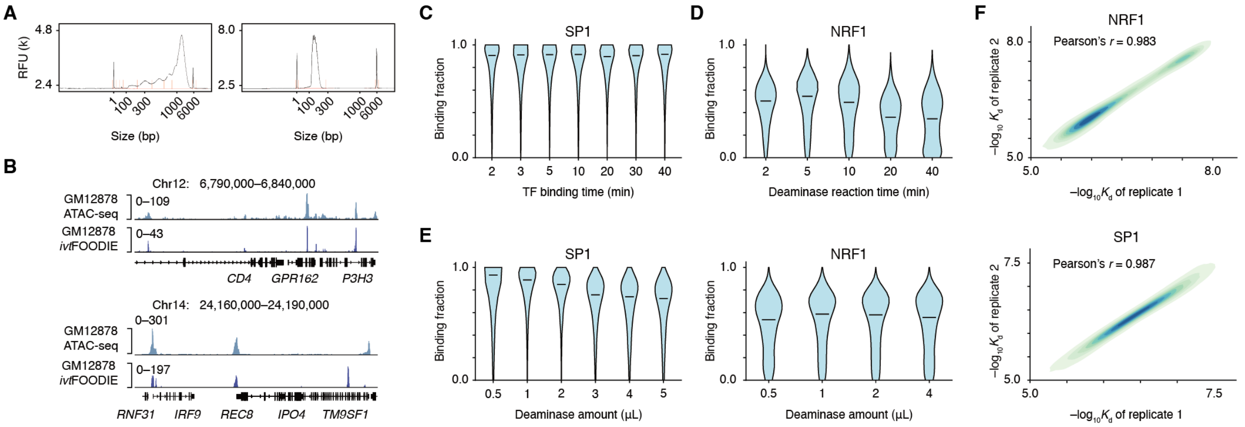


**Fig. S1. Optimization and validation of the *ivt*FOODIE assay conditions. (A)** Capillary electrophoresis profiles of GM12878 open chromatin DNA fragments before enrichment (left) and after size selection by gel extraction (right). **(B)** Representative genome browser tracks comparing ATAC-seq and signals from *ivt*FOODIE input libraries across selected loci in GM12878 cells. Annotated gene structures are shown at the bottom. **(C)** Distribution of SP1 binding fractions across GM12878 open chromatin regions at various TF binding times. **(D)** Distribution of NRF1 binding fractions across the open chromatin regions following different deaminase reaction times. **(E)** Effects of varying deaminase amounts on the binding fractions of SP1 (left) and NRF1 (right). **(F)** Reproducibility of the *ivt*FOODIE assay. Scatter plots display the correlation of estimated $-\log_{10}K_d$ values between two independent replicates for NRF1 (top) and SP1 (bottom).

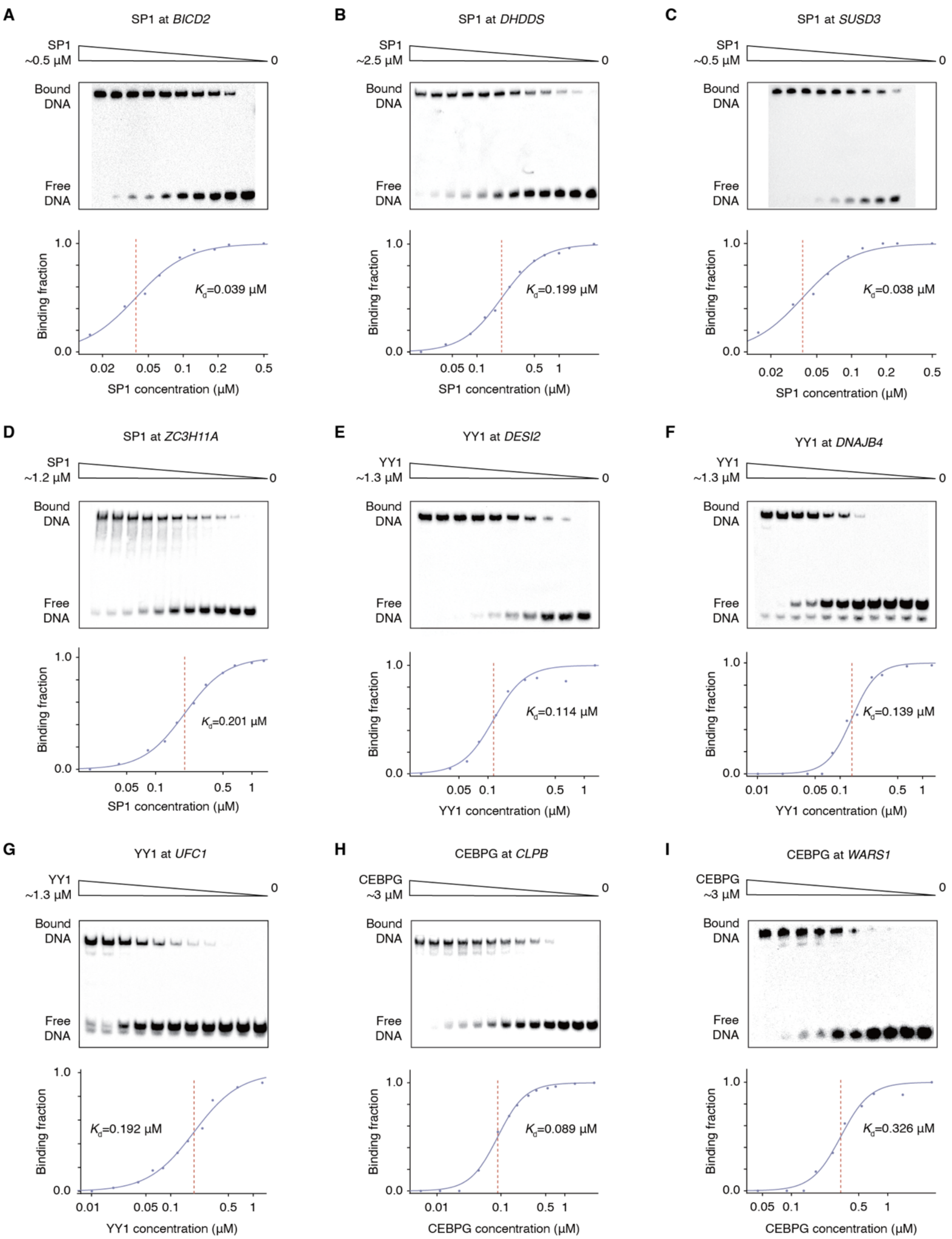


**Fig. S2. Quantification of SP1, YY1, and CEBPG binding affinities by EMSA. (A–D)**, SP1 binding to the regulatory elements of *BICD2*, *DHDDS*, *SUSD3*, and *ZC3H11A*. Top: Representative EMSA gel images. Bottom: Quantification of binding fractions based on DNA band intensities. Protein concentrations are plotted on a logarithmic scale. Purple curves indicate fitting using the Hill equation, and orange dotted lines mark the calculated dissociation constants. **(E–G)**, YY1 binding to the regulatory elements of *DESI2*, *DNAJB4*, and *UFC1*, analyzed and presented as in **(A–D)**. **(H–I)**, CEBPG binding to the regulatory elements of *CLPB* and *WARS1*, analyzed and presented as in **(A–D)**. For all panels, data shown are representative of three independent experiments with consistent results.

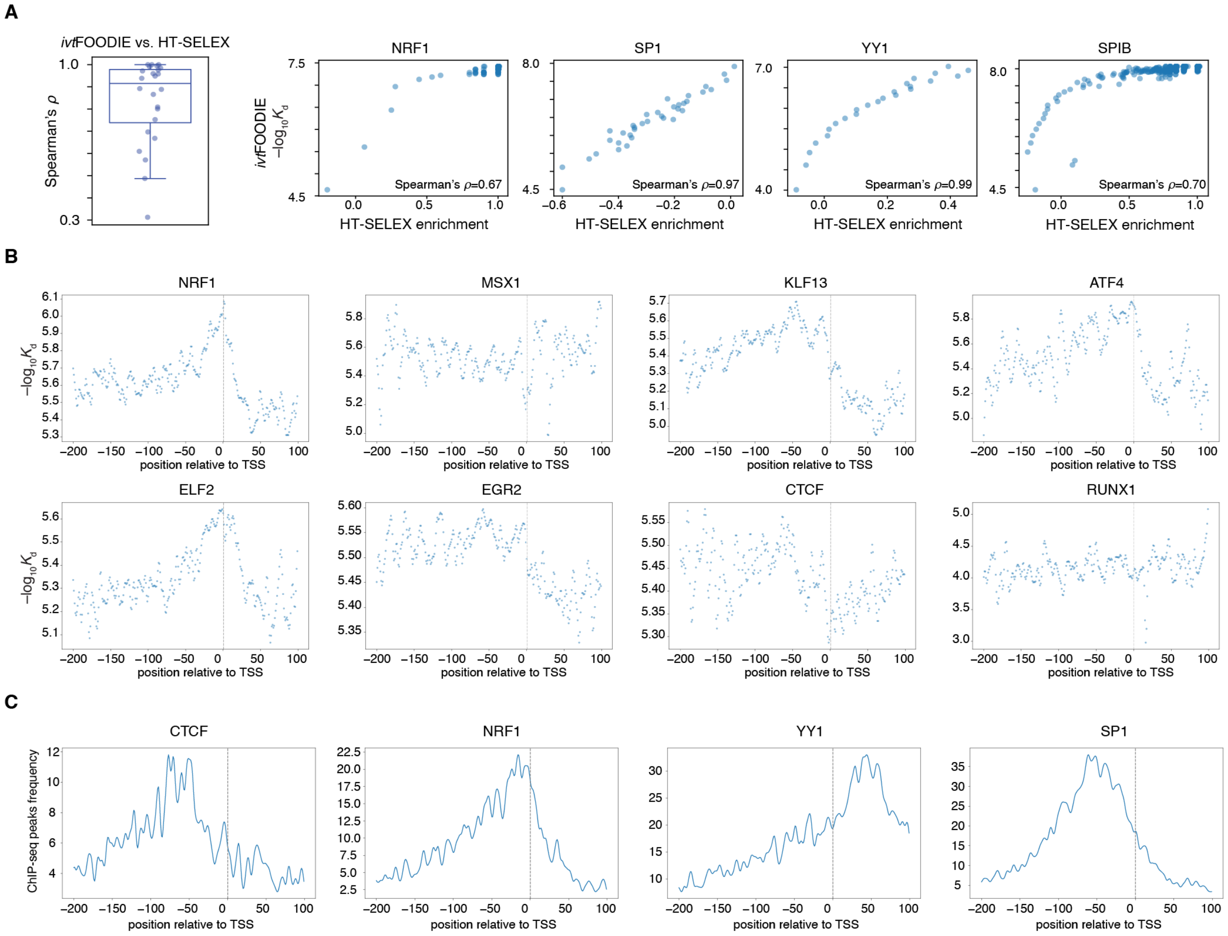


**Fig. S3. Validation of *ivt*FOODIE predictions and spatial profiling of TF binding relative to TSSs. (A)** Concordance between ivtFOODIE and HT-SELEX. The boxplot (left) displays the distribution of Spearman's $\rho$ between experimental *ivt*FOODIE $-\log_{10}K_d$ values and HT-SELEX enrichment scores across all evaluated TFs. Scatter plots (right) further illustrate this positive correlation for four representative TFs. **(B)** Profiles of TF binding affinities near TSSs. Scatter plots represent the mean *ivt*FOODIE $-\log_{10}K_d$ values aggregated across the TSSs (position 0; vertical dashed lines) of all protein-coding genes for eight representative TFs. **(C)** Spatial frequency of *in vivo* TF binding events. Line plots depict the cumulative count of ChIP-seq peak summits mapped relative to the TSS for four representative TFs.

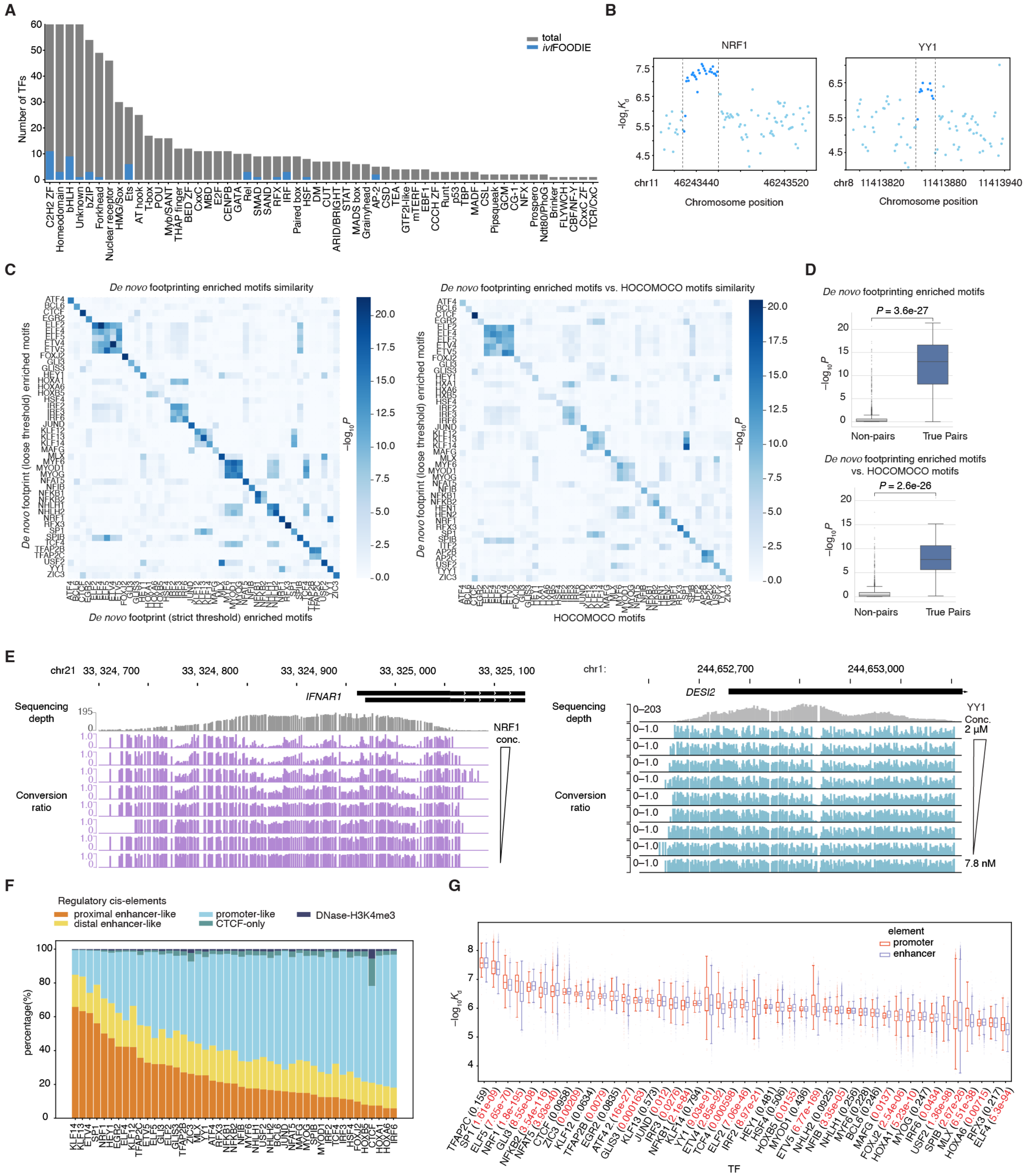


**Fig. S4. The landscape and binding affinities of TF binding sites. (A)** Distribution of the number of TFs across distinct structural families in the human genome. TFs characterized in the *ivt*FOODIE experiments are highlighted in blue. **(B)** Single-cytosine resolution $–\log_{10}K_d$ values mapped across specific genomic loci for representative TFs, NRF1 and YY1. Vertical gray dashed lines indicate the boundaries of *de novo*-identified footprints. **(C)** Heatmaps depicting motif similarity scores. Left: Comparison of *de novo* footprint-enriched motifs identified using strict versus loose threshold parameters. Right: Comparison between *de novo* footprint-enriched motifs (loose threshold) and reference motifs from the HOCOMOCO database. Similarity scores ($–\log_{10}P$)

were calculated using Tomtom. **(D)** Boxplots comparing the similarity scores of paired versus non-paired motifs, corresponding to the heatmaps in **(C)**. Paired motifs represent comparisons for the same TF (either between de novo motifs identified under different footprinting thresholds, or between *de novo* and HOCOMOCO reference motifs). Non-paired motifs represent comparisons between different TFs. **(E)** Representative *ivt*FOODIE genome browser tracks at the *IFNAR1* (left) and *DESI2* (right) loci. Tracks display sequencing depth and cytosine conversion ratios across a decreasing titration gradient of NRF1 and YY1 concentrations. **(F)** Relative proportion of distinct regulatory *cis*-elements intersecting with *de novo* identified footprints across the indicated TFs. **(G)** Comparison of calculated TF $-\log_{10}K_d$ values within promoter (red) versus enhancer (blue) regions for various TFs. *P* values are indicated in parentheses on the x-axis, with significant values ($P < 0.05$) highlighted in red.

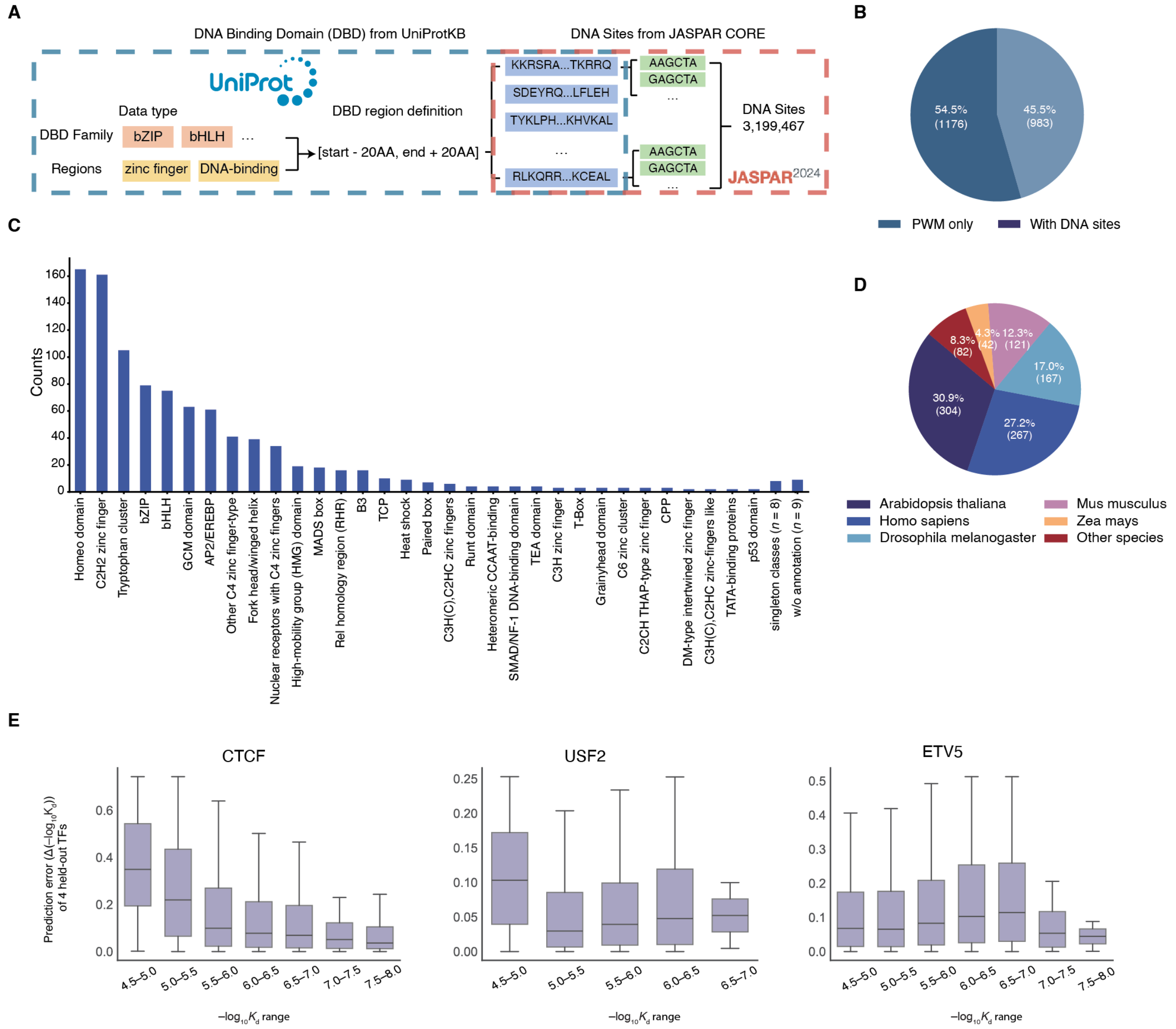


**Fig. S5. Pre-training data curation workflow and summary statistics. (A)** Workflow for assembling TF sequences with associated DNA-binding sites. The annotated DBD boundaries were symmetrically extended by 20 amino acids (AA) at both termini (start − 20 AA, end + 20 AA). Experimentally validated DNA-binding sites were curated from the JASPAR CORE database (2024 release; $n$ = 3,199,467 sites). The schematic illustrates representative TF subsequences and their matched DNA-binding sites. **(B)** Coverage of TFs in JASPAR. Proportions of TFs with only PWM data available (54.5%, $n$ = 1,176) versus TFs with both PWMs and experimentally determined DNA-bound sites (45.5%, $n$ = 983). **(C)** Distribution of TF DBD families in the resource. Homeodomain and C2H2 zinc-finger families are most frequent, followed by tryptophan-cluster, bZIP, bHLH, GCM, AP2/ERF, forkhead/winged-helix, and other families; bars indicate counts per family. **(D)** Species composition of TFs with experimentally determined DNA-bound sites (n = 983). The largest contributions are from Arabidopsis thaliana (30.9%; n = 304) and Homo sapiens (27.2%; n = 267), followed by Zea mays (17.0%; n = 167), Mus musculus (12.3%; n = 121), Drosophila melanogaster (8.3%; n = 82), and other species (4.3%; n = 42). **(E)** Relationship between model prediction error and binding affinity for representative TFs. Box plots display the prediction error ($\Delta(-\log_{10}K_d)$) across varying ranges of $-\log_{10}K_d$ values for three representative held-out TFs: CTCF, USF2, and ETV5.

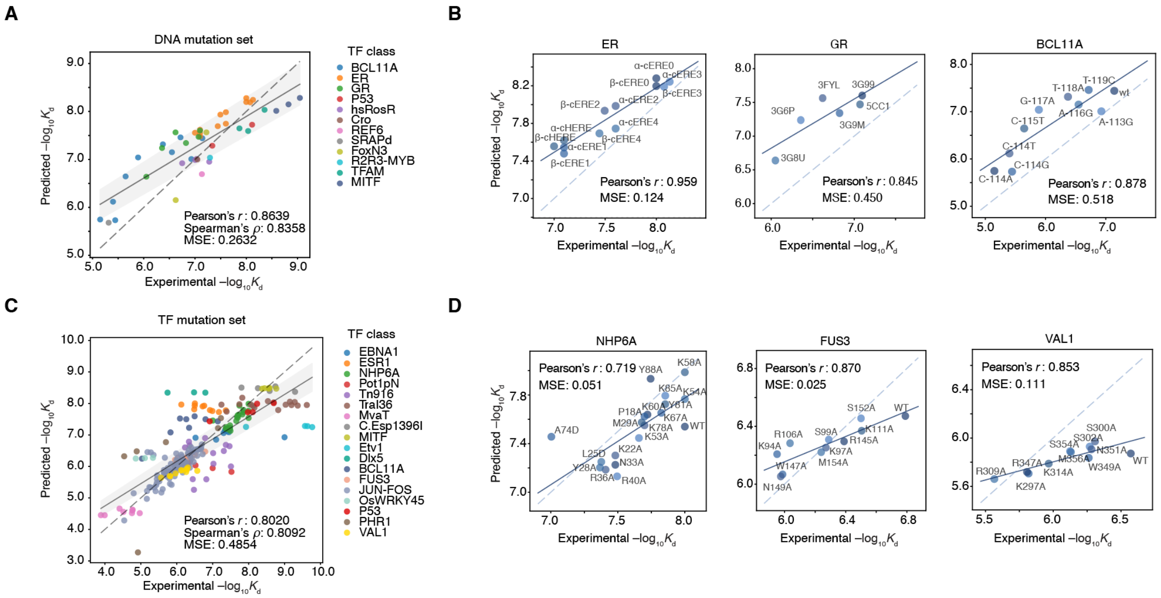


**Fig. S6. Benchmarking predicted TF–DNA affinities against experimental data. (A)** Overall results of DNA-mutation datasets. Each category represents the binding affinities of the same TF paired with multiple different DNA sequences. **(B)** Scatter plots for the ER, GR and BCL11A datasets with different DNA sequences. **(C)** Overall results of TF-mutation datasets. The scatter plot displays the correlation between predicted and experimentally measured TF–DNA binding affinities for cases involving TF mutations. Each point corresponds to a unique TF–DNA pair, with different TF categories denoted by distinct colors. Each category includes the binding affinities of mutant TFs paired with the same DNA sequence. The x-axis represents the experimentally measured affinity values ($–\log_{10}K_d$) and the y-axis shows the corresponding predicted affinity values ($–\log_{10}K_d$). **(D)** Scatter plot on NHP6A, FUS3 and VAL1 datasets with mutant TF sequences, providing detailed information such as mutant positions, correlation coefficients and MSE values between predicted and experimental affinities for these particular cases.

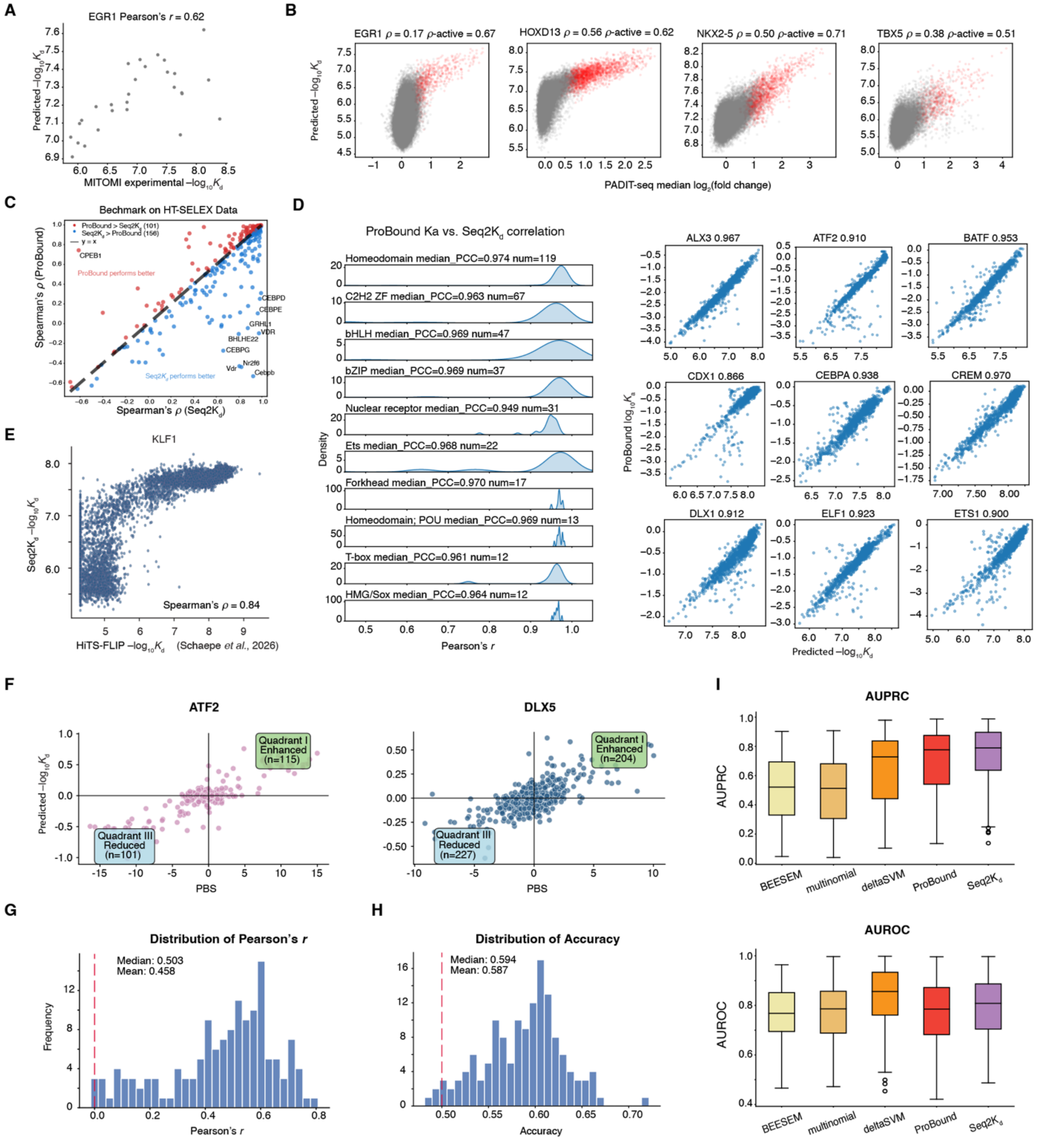


**Fig. S7. Benchmarking the performance of Seq2K$_d$. (A)** Correlation between Seq2K$_d$-predicted and MITOMI-measured binding affinities for EGR1. **(B)** Comparison of Seq2K$_d$-predicted and PADIT-seq-measured affinities for EGR1, HOXD13, NKX2-5, and TBX5. Red dots represent DNA sequences identified by PADIT-seq as "active" (supporting *in vitro* transcription) while gray dots are inactive sequences. $\rho$ and $\rho$-active denote Spearman's correlation coefficients computed for all sequences and the active sequence subset, respectively. **(C)** Performance comparison of Seq2K$_d$ and ProBound on HT-SELEX data. The scatter plot displays Spearman's $\rho$ between predicted affinities and experimental enrichment fold changes for each respective model. **(D)** Distributions of Pearson's $r$ between Seq2K$_d$- and ProBound-predicted affinities across diverse

DBD classes (left). Representative scatter plots for individual TFs are shown on the right. **(E)** Correlation between Seq2$K_d$-predicted and HiTS-FLIP-derived experimental $-\log_{10}K_d$ values for KLF1. **(F)** Scatter plots showing predicted $-\log_{10}K_d$ values versus PBS (preferential binding score) for two representative TFs (ATF2 and DLX5) from the novel batch of SNP-SELEX experiments. SNPs in the upper-right or lower-left quadrants correspond to enhanced or reduced binding. **(G)** The distribution of Pearson's *r* values across TFs for Seq2$K_d$ predcitions on the novel batch of SNP-SELEX experiments. **(H)** The distribution of accuracy (the proportion of correctly predicted directions of affinity changes between two alleles of the SNP) across TFs between Seq2$K_d$ and the novel batch of SNP-SELEX experiments. **(I)** SNP-SELEX benchmark. Comparison of AUPRC and AUROC in predicting pbSNPs in the novel SNP-SELEX batch among Seq2$K_d$, ΔPWM-BEESEM, ΔPWM-multinomial, deltaSVM, and ProBound.

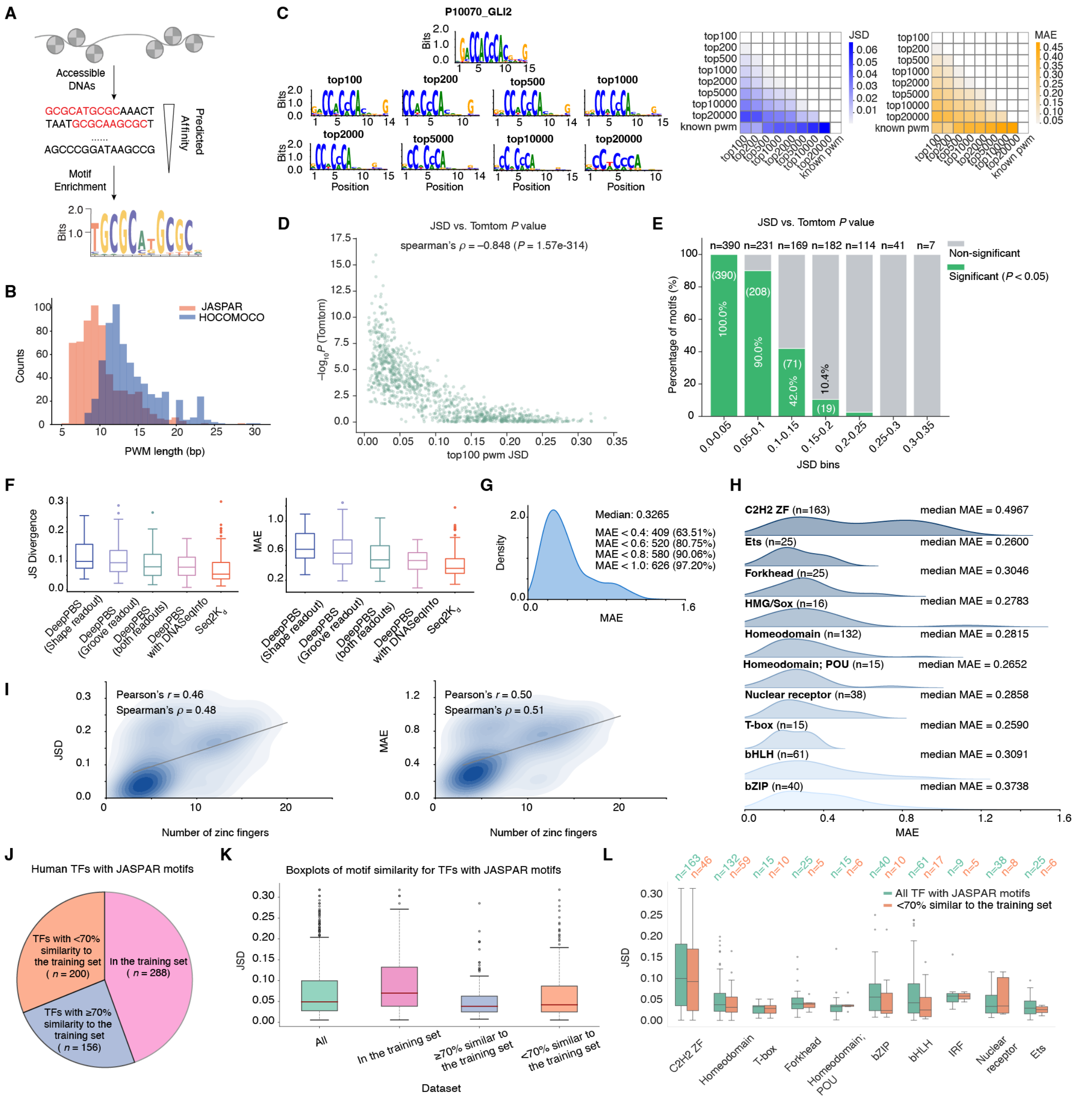


**Fig. S8. Workflow and validation of *de novo* motif derivation via Seq2K$_d$. (A)** Schematic of the motif discovery workflow, where motifs (PWM logos) are derived from accessible DNA fragments ranked by Seq2K$_d$-predicted affinities. **(B)** Distribution of PWM lengths in JASPAR and HOCOMOCO databases. **(C)** Comparison between JASPAR PWM and Seq2K$_d$-derived PWMs for GLI2. Left, PWM logos enriched from sequences across various top-K predicted affinity thresholds (K = 100 to 20,000). Including more sequences resulted in minimal changes in the PWMs. Right, heatmaps showing pairwise JSD (top) and MAE (bottom) between the reference and predicted PWMs at different K values. **(D)** Correlation between JSD and Tomtom significance. Scatter plot showing the relationship between JSD values and Tomtom $-\log_{10}P$ values. **(E)** Proportion of significant motif matches across JSD intervals. The percentage of predicted motifs categorized as significant (Tomtom $P < 0.05$, green) or non-significant (gray) across increasing JSD intervals. **(F)** Performance comparison between Seq2K$_d$ and four DeepPBS variants. Boxplots show the distribution of JSD (left) and MAE (right) between predicted and experimental PWMs. **(G)** Kernel density distribution of global MAE values between Seq2$K_d$-predicted motifs and JASPAR references across all evaluated TFs. **(H)** MAE density distributions stratified by TF family. The median MAE and number of TFs (*n*) are annotated for each family. **(I)** Accuracy of C2H2-ZF motif predictions relative to finger count. Kernel density estimation (KDE) plots showing the number of C2H2 zinc fingers versus JSD (left) and MAE (right) between JASPAR reference and predicted motifs. **(J)** Homology-depleted held-out set construction. Partitioning of JASPAR human TFs into the training set ($n$ = 288), TFs with ≥70% DBD similarity to training samples ($n$ = 156), and a strictly independent held-out set (<70% similarity, $n$ = 200). **(K)** Seq2$K_d$ performance across partitions. JSD distributions for the total human TF collection ($n$ = 644) and the three subsets defined in **(J)**. **(L)** Performance across TF families. Comparison of JSD values between all JASPAR human TFs (green) and the independent held-out set (orange), stratified by major TF families. Sample sizes (*n*) for each group are indicated at the top.

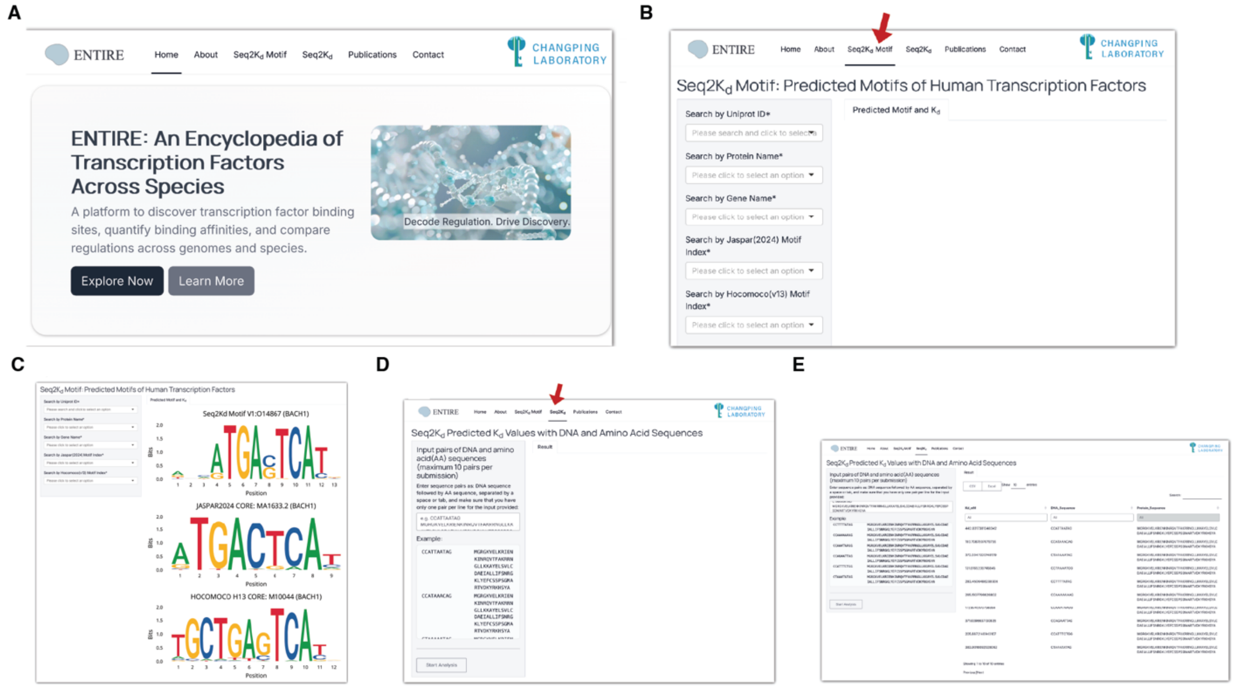


**Fig. S9. The ENTIRE web portal for human TF motif and Seq2K$_{\mathrm{d}}$ prediction. (A)** Overview of the ENTIRE web portal (https://www.entire.ac.cn/). The platform comprises two primary modules: the Seq2K$_{\mathrm{d}}$ Motif (V1) database and the Seq2K$_{\mathrm{d}}$ tool. The database extends existing JASPAR and HOCOMOCO resources by providing enhanced, predicted human TF motifs, notably covering proteins lacking reference motifs in these public databases. Concurrently, the Seq2K$_{\mathrm{d}}$ tool predicts the $K_{\mathrm{d}}$ values for user-submitted AA and DNA sequence pairs. The landing page offers direct access to both modules alongside supplementary portal information. **(B)** The Seq2K$_{\mathrm{d}}$ Motif (V1) database interface. Users can query human TFs by UniProt ID, protein name, gene symbol, or existing motif indices from JASPAR or HOCOMOCO. Search results yield predicted motifs and $K_{\mathrm{d}}$ values, summarized in an integrated table. **(C)** An example query for the BACH1 protein. The panel displays the Seq2K$_{\mathrm{d}}$ Motif (V1) predicted motif (top) aligned with reference motifs from JASPAR and HOCOMOCO. Sequence logos are represented as information content (bits) per position, facilitating a direct visual comparison of motif features. **(D)** The Seq2K$_{\mathrm{d}}$ tool interface for $K_{\mathrm{d}}$ prediction. Users can submit up to 10 protein–DNA pairs per job, formatting each input as an AA sequence and a DNA sequence separated by a space. **(E)** Example analysis output displaying the results for submitted sequence pairs. The table reports the predicted $K_{\mathrm{d}}$ values alongside the corresponding input DNA and AA sequences. The interface provides search, pagination, and row-adjustment functionalities to facilitate efficient data review.

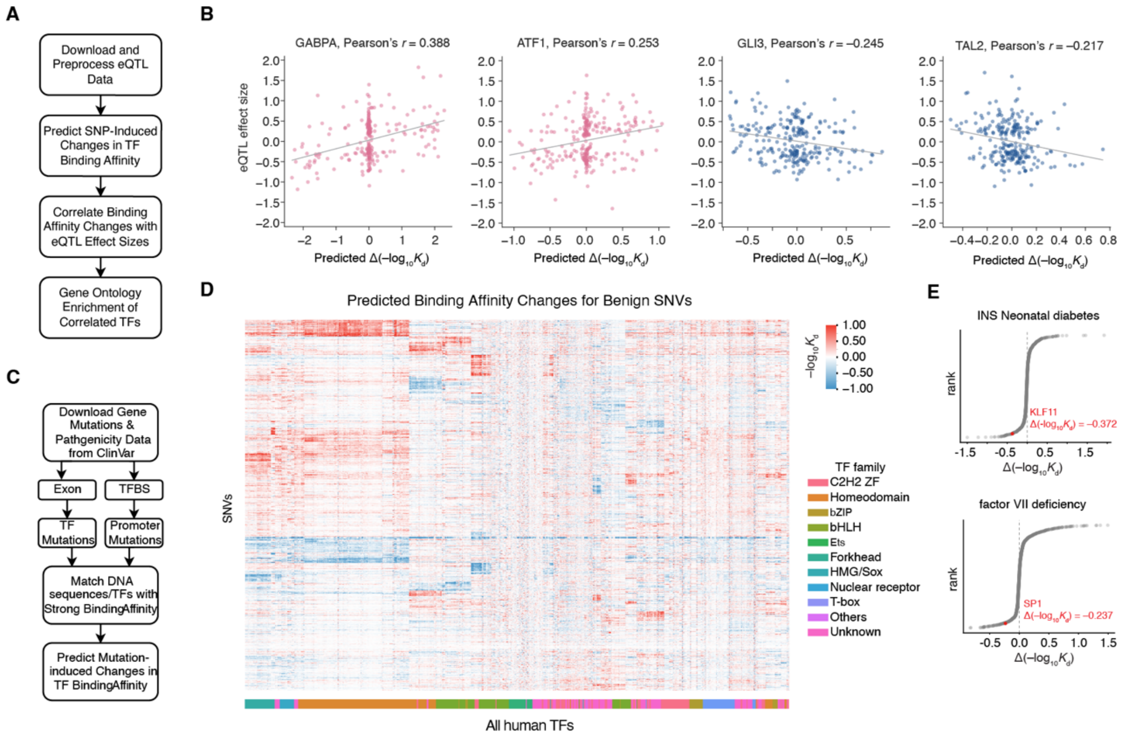


**Fig. S10. Evaluation of Seq2K$_d$ in predicting the functional impact of non-coding variants on TF binding and regulatory activity. (A)** Workflow for correlating eQTL effect sizes with predicted changes in TF binding affinities. **(B)** Correlations between Seq2K$_d$-predicted binding affinity changes and eQTL effect sizes. Data points (representing individual eQTLs) are colored red or blue for TFs that exhibit overall positive or negative correlations, respectively. **(C)** Workflow for evaluating the impact of benign and pathogenic SNVs in TF DBD-coding regions and promoter non-coding regions on TF–DNA binding affinities. **(D)** Heatmap of predicted binding affinity changes ($\Delta(-\log_{10}K_d)$) across all human TFs (columns) for benign promoter SNVs curated from ClinVar (rows). **(E)** Scatter plots of predicted affinity changes versus rank for all human TFs at the mutant promoter regions of *INS* (neonatal diabetes) and *F7* (factor VII deficiency). Red dots highlight the specific affected TFs (KLF11 and SP1, respectively).

**table S1.**
Summary of TF expression constructs, purification tags, and deaminase sequences used in the *ivt*FOODIE assay.

**table S2.**
Oligonucleotide sequences used in EMSA to validate TF–DNA binding affinities.
